# Glycolytic enzymes localize to synapses under energy stress to support synaptic function

**DOI:** 10.1101/042002

**Authors:** SoRi Jang, Jessica C. Nelson, Eric G. Bend, Lucelenie Rodríguez-Laureano, Felipe G. Tueros, Luis Cartagenova, Katherine Underwood, Erik M. Jorgensen, Daniel A. Colón-Ramos

**Author notes:** Correspondence to: Daniel A. Colón-Ramos, Ph.D. Department of Cell Biology, Yale Program in Cellular Neurodegeneration and Repair, Yale University School of Medicine, 295 Congress Avenue, BCMM 436B, New Haven, CT 06510. These authors contributed equally to this work (SoRi Jang, Jessica C. Nelson).

## Abstract

Changes in neuronal activity create local and transient changes in energy demands at synapses. Here we discover a metabolic compartment that forms *in vivo* near synapses to meet local energy demands and support synaptic function in *Caenorhabditis elegans* neurons. Under conditions of energy stress, glycolytic enzymes redistribute from a diffuse localization in the cytoplasm to a punctate localization adjacent to synapses. Glycolytic enzymes colocalize, suggesting the *ad hoc* formation of a glycolysis compartment, or a ‘glycolytic metabolon’, that can maintain local levels of ATP. Local formation of the glycolytic metabolon is dependent on presynaptic scaffolding proteins, and disruption of the glycolytic metabolon blocks the synaptic vesicle cycle, impairs synaptic recovery, and affects locomotion. Our studies indicate that energy demands in neurons are met locally through the assembly of a glycolytic metabolon to sustain synaptic function and behavior.

## Introduction

The brain consumes more energy than any other organ in the body, accounting for ~20% of total body energy consumption (Belanger et al., 2011; Mink et al., 1981). Within the brain, synapses are primary sites of ATP consumption—the synaptic vesicle cycle being one of the main sources of activity-driven metabolic demands (Harris et al., 2012; Rangaraju et al., 2014). Conditions that alter the metabolic state of the brain, such as hypoxia, starvation, and hypoglycemia, have profound effects on synaptic transmission and cognitive function (Cherubini et al., 1989; Gold et al., 1995). Even brief interruptions of activity-stimulated ATP synthesis can result in severe impairment of synaptic function (Rangaraju et al., 2014).

ATP is predominantly produced by either glycolysis or oxidative phosphorylation. But not all ATP is created equal, and these two sources contribute differentially to various metabolic processes (Pfeiffer et al., 2001). Oxidative phosphorylation, which is mediated by the mitochondria, is an efficient process that produces high yields of ATP molecules, but at low rates of production. Glycolysis, on the other hand, can act independently from the mitochondria to produce lower yields of ATP, but at faster rates (Pfeiffer et al., 2001). Tissues that consume ATP at higher rates, including the brain (but also muscles, developmental tissues, and cancer cells) heavily rely on the glycolytic machinery to meet their energy demands (Vander Heiden et al., 2009; Wojtas et al., 1997). While glycolysis is inhibited by oxygen in most cells, in these tissues glycolysis is active under aerobic conditions. The preferential use of glycolysis over oxidative phosphorylation even in aerobic conditions is referred to as the *Warburg effect*, or aerobic glycolysis (Warburg et al., 1927). Aerobic glycolysis plays an important role in brain metabolism and function (Gjedde and Marrett, 2001; Magistretti and Allaman, 2013) and increases locally upon conditions of increased neuronal activity (Vaishnavi et al., 2010).

Neuronal activity can change the energy demands at synapses (Harris et al., 2012; Rangaraju et al., 2014). Because the diffusion rate of intracellular ATP is limited (Hubley et al., 1996), synapses must rely on local production of ATP to meet these transient changes in energy demands and sustain synaptic function. Mitochondria, which mediate oxidative phosphorylation, are actively transported to neuronal synapses, and defects in localization have been linked to neurodegenerative disorders such as Parkinson’s disease (Burte et al., 2015; Lin and Sheng, 2015; Schwarz, 2013). Unlike the mitochondrion, which is a membrane-bound organelle, glycolytic enzymes are soluble proteins in the cytosol. Glycolytic enzymes, however, are not uniformly distributed throughout the cytosol (Masters, 1991; Menard et al., 2014), and in neurons, biochemical studies demonstrated that glycolytic proteins are enriched in synaptic fractions (Knull, 1978, 1980; Knull and Fillmore, 1985). The physiological importance of the localization of glycolytic enzymes, and their role in meeting local energy demands at synapses remains poorly understood.

In this study we identify, from forward genetic screens in *C. elegans*, a role for the glycolytic machinery in powering the synaptic vesicle cycle. We demonstrate that under conditions of energy stress, glycolytic proteins dynamically colocalize near presynaptic sites into a metabolic compartment. We also demonstrate that presynaptic scaffolding proteins are necessary for the *ad hoc* localization of glycolytic proteins to presynaptic sites, and that the local assembly of this metabolic compartment is necessary for synaptic function and locomotion under energy stress. Our studies indicate that energy demands in *C. elegans* neurons are met locally through the assembly of a glycolytic metabolon to sustain synaptic function and behavior.

## Results

### Glycolytic proteins are required to maintain synaptic vesicle clusters during hypoxia

We conducted unbiased forward genetic screens to identify molecules required for the localization of synaptic vesicle proteins in the serotonergic NSM neurons in the nematode *C. elegans* (Figure 1A-1D, 1H; genetic screen described in Figure S1 and *Experimental Procedures*). From this screen we identified allele *ola72*, which displayed diffuse distribution of the synaptic vesicle proteins VMAT/CAT-1 and synaptobrevin under hypoxic conditions (Figure 1E-1G, 1I, and 1N). Positional cloning of the *ola72* allele revealed a missense mutation (C562Y) in *pfk-1.1* (Figure S2A) – one of two *C. elegans* genes that encode phosphofructokinase-1. In mutant animals carrying the *ola72* allele, synaptic vesicle proteins cluster normally under normoxic conditions (Figures 1J, 1M, 1O, and S1C). However, under hypoxic conditions (induced by mounting animals under a glass coverslip (Pitts and Toombs, 2004), or by incubation in a hypoxia chamber) these synaptic vesicle proteins become diffusely distributed throughout the neurite in *ola72* animals (Figures 1J, 1N, 1P, and S1F; Movie S1). This phenotype was not observed in wild-type animals, which maintained punctate localization of synaptic vesicle proteins under the same hypoxic conditions (Figures 1J-1L, and S1E). Three independent alleles, *pfk-1.1(gk549413), pfk-1.1(gk758818)*, and *pfk-1.1(gk922689)*, phenocopy and fail to complement the *ola72* allele (Figures 2C, 2I, and S2C). Expression of a wild-type copy of the *pfk-1.1* gene in *ola72* mutant animals rescues punctate localization of synaptic vesicle proteins under hypoxic conditions (Figure S2B). These findings indicate that phosphofructokinase-1 is required to maintain the localization of synaptic vesicle proteins at synapses under hypoxic conditions.

**Figure 1.**
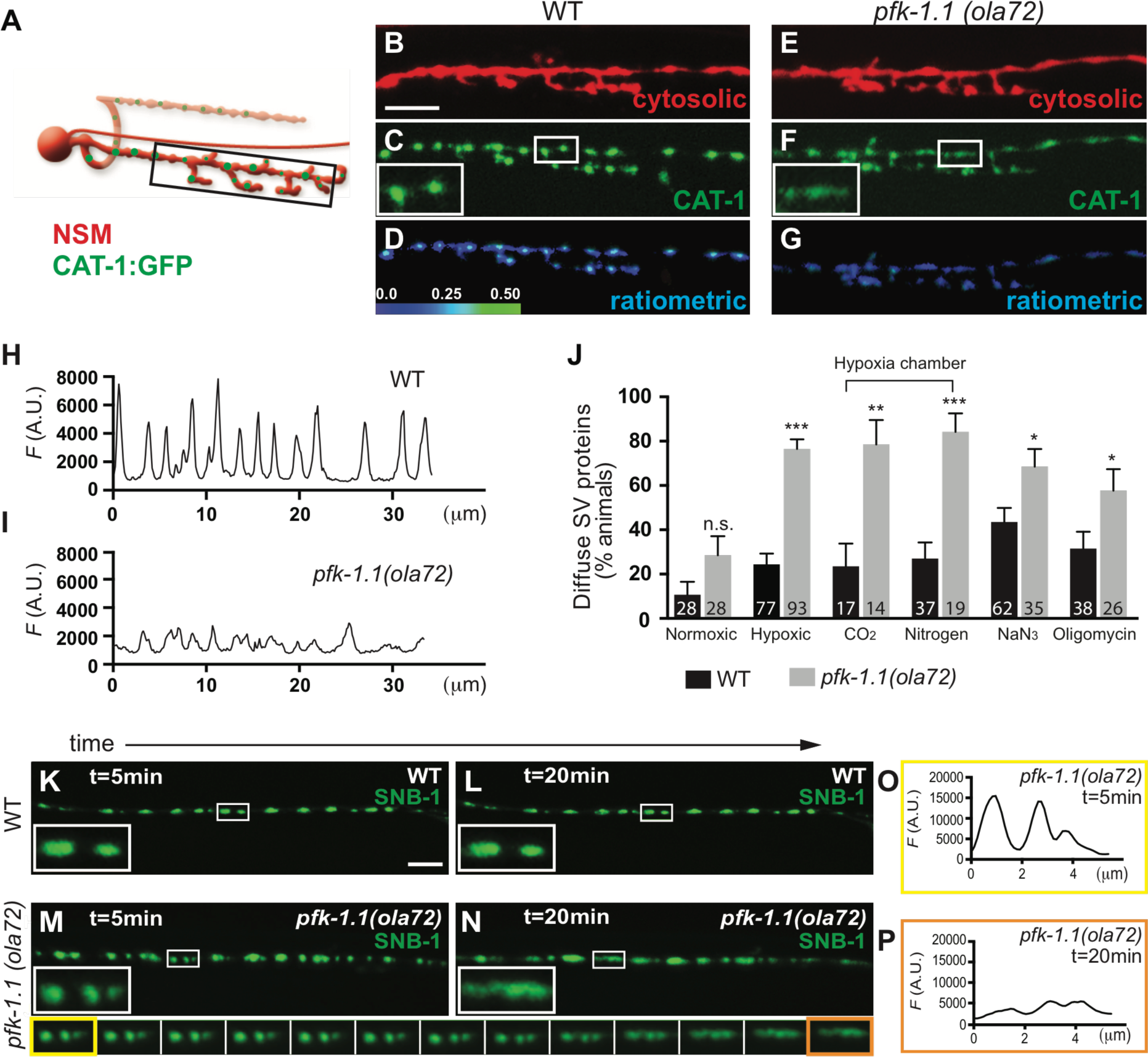
PFK-1.1 is required to cluster synaptic vesicle proteins under hypoxic conditions. (A) Schematic of NSM neuron (red) with synaptic release sites (green), as previously described (Axang et al., 2008; Nelson and Colón-Ramos, 2013). Black box marks the ventral neurite imaged in panels B-G. (B-G) The ventral neurite of the serotonergic NSM neurons (B and E) and its synaptic vesicle proteins (GFP tagged vesicular monoamine transporter VMAT/CAT-1) were simultaneously visualized in wild-type (B-C) and *pfk-1.1(ola72)* mutant animals (E-F) under hypoxic conditions (hypoxia was induced as described in *Experimental Procedures*). Ratiometric images of NSM represent the relative enrichment of CAT-1::GFP signal over cytosolic mCherry signal for wild-type (D) or *pfk-1.1(ola72)* mutant animals (G). (H-I) Pixel fluorescence values along the ventral neurite of the wild-type (H) and *pfk-1.1 (ola72)* (I), corresponding to images (C) and (F), respectively. (J) Percentage of animals displaying a diffuse distribution of synaptic vesicle proteins in wild-type or *pfk-1.1 (ola72)* mutant animals under varying conditions. Hypoxic conditions were induced by mounting animals on a glass slide under a glass coverslip for 10 minutes (Pitts and Toombs, 2004), or in a hypoxia chamber for 10 minutes, or 10 minutes of pharmacological treatment, as described in the *Experimental Procedures*. Wild-type and *pfk-1.1(ola72)* mutant animals were also scored blindly (see *Experimental Procedures*). Number of animals scored is indicated at the bottom of each column. (K-N) Time-lapse displaying the distribution of synaptic vesicle proteins in GABA neurons (imaged with synaptobrevin/SNB-1::GFP) of a single wild-type (K, L); or *pfk-1.1 (ola72)* mutant worm (M, N); after 5 minutes (K, M); or 20 minutes (L, N) under hypoxic conditions. Insets correspond to zoomed-in (3×) images of the indicated regions. Bottom row corresponds to time-lapse images (1 minute intervals) of the insets in *pfk-1.1(ola72)* (Please also see Movie S1, a time-lapse movie of synaptic vesicles in *pfk-1.1 (ola72)*). Note vesicular markers remain clustered in wild-type animals (L), but become diffusely distributed in *pfk-1.1 (ola72)* mutant animals over time under hypoxia (N and bottom row). (O-P) Line scan pixel fluorescence values for the first inset in the montage (O) (M, also yellow trim in bottom row) and the last inset (P) (N, also orange trim in bottom row). In all panel, scale bar represents 5μm. Error bars denote s.e.m. *, p < 0.05. **, p < 0.01. ***, p < 0.001 as compared to wild-type animals under similar conditions.

**Figure 2.**
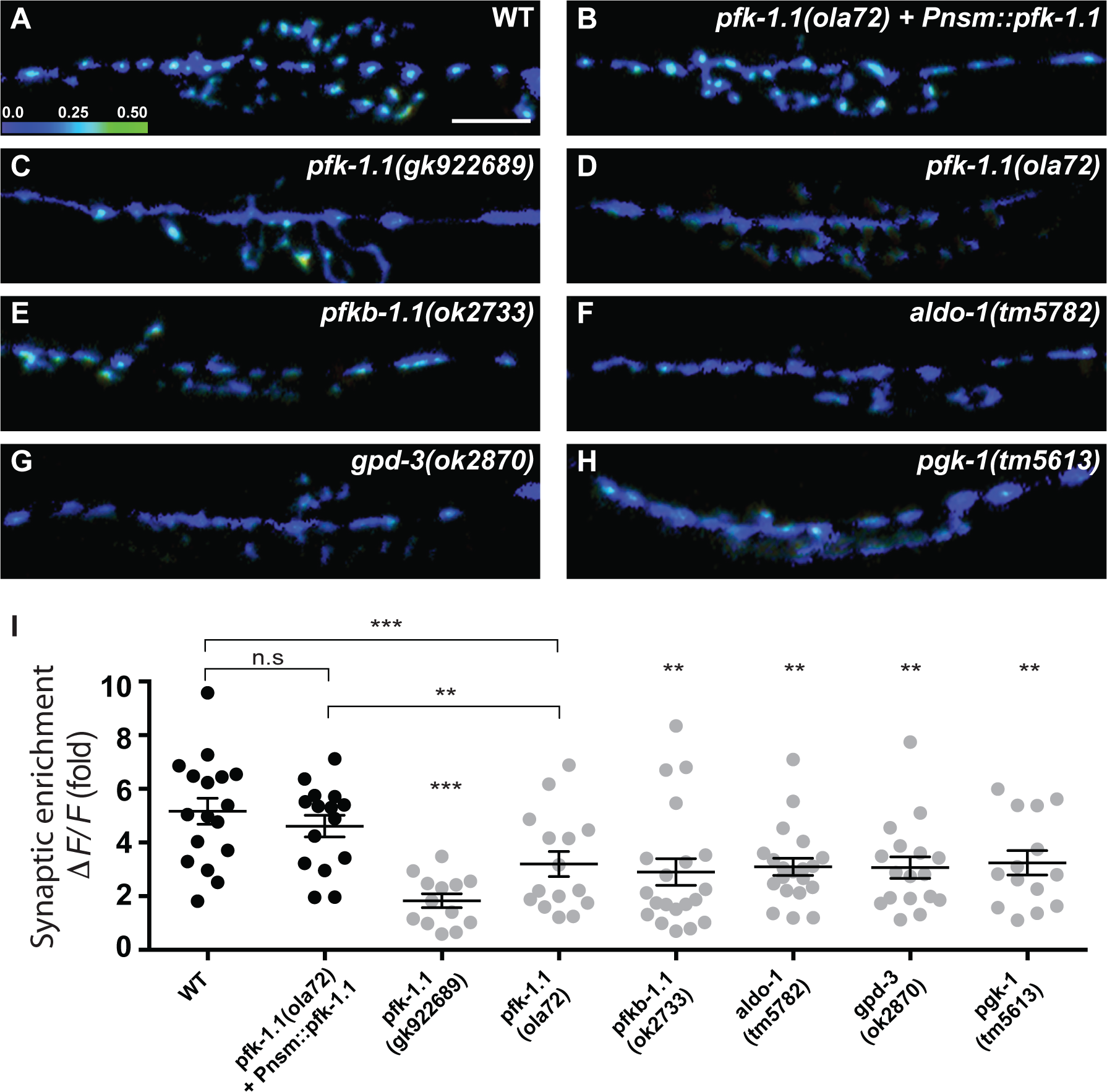
Glycolytic proteins are required to maintain synaptic vesicle clusters during hypoxia. (A-H) Ratiometric images of NSM representing the relative enrichment of CAT-1::GFP signal over cytosolic mCherry signal for wild-type (A); *pfk-1.1(ola72)* mutant animals expressing a wild-type rescuing array of *pfk-1.1* cell-autonomously in NSM (B); *pfk-1.1(gk922689)* (C); *pfk-1.1(ola72)* (D); *pfkb-1.1(ok2733)* (E); *aldo-1(tm5782)* (F); *gpd-3(ok2870)* (G); and *pgk-1(tm5613)* (H). Note that *pfk-1.1(gk922689)* phenocopies the *ola72* allele under hypoxic conditions, indicating that, along with positional mapping, complementation tests and rescue experiments, *ola72* is an allele of *pfk-1.1* (see also Figures S2A-S2C). Note also that expression of a wild-type copy of *pfk-1.1* cDNA cell-specifically in the NSM neurons rescues the diffuse distribution of synaptic vesicle proteins in NSM (B), indicating the *pfk-1.1* acts cell autonomously. (I) Quantification of the distribution of the synaptic vesicle proteins for the examined genotypes. Percent change between peak and trough fluorescence values, or synaptic enrichment of the synaptic vesicle proteins, (*ΔF/F*) in the NSM ventral neurite was calculated as described (Dittman and Kaplan, 2006) (See also *Experimental Procedures*). The circles in the graph represent individual animals. Scale bar in (A) represents 5μm and is applicable to all panels. Error bars denote s.e.m. *, p < 0.05. **, p < 0.01. ***, p < 0.001 as compared to wild-type animals under similar conditions, unless otherwise indicated by brackets.

The *pfk-1.1* phenotype is not due to a disruption of the synapses themselves, since the localization of the presynaptic active zone protein ELKS is not altered in *pfk-1.1* mutants (Figures S3A-S3E). In addition, the requirement for *pfk-1.1* is not limited to the NSM neuron, but is also required for the maintenance of vesicle protein clustering in all neurons examined, including the nerve ring interneuron AIY, GABA motor neurons, and acetylcholine motor neurons (Figures 1K-1P, S3A-S3J). Consistent with the pan-neuronal phenotype, a GFP reporter under the *pfk-1.1* promoter is expressed in all examined neurons, including the NSM neuron (Figures S2D-S2G). Moreover, NSM neuron-specific expression of the *pfk-1.1* cDNA in *pfk-1.1(ola72)* mutants led to rescue of the synaptic vesicle phenotype in the NSM neuron (Figures 2B and 2I). Our findings indicate that the phosphofructokinase enzyme is required cell autonomously in neurons to cluster synaptic vesicle proteins under hypoxic conditions.

Phosphofructokinase-1 is a rate-limiting enzyme that catalyzes the first committed step during glycolysis. To determine if the glycolytic pathway is required for the maintenance of vesicle clusters in neurons under hypoxic conditions, we examined mutants of other glycolytic enzymes (Figure S2H). Mutants for phosphofructokinase-2/fructose-2,6-bisphosphatase/*pfkb-1.1(ok2733)*, glyceraldehyde 3-phosphate dehydrogenase*/gpd-3(ok2870)*, aldolase/*aldo-1(tm5782)*, and phosphoglycerate kinase*/pgk-1(tm5613)* phenocopy the hypoxia-dependent synaptic vesicle phenotype of *pfk-1.1* mutant animals (Figures 2C-2I). To examine if the observed role of glycolysis in the maintenance of vesicle clusters during hypoxia was due to disruption of oxidative phosphorylation, we pharmacologically blocked cytochrome oxidase with sodium azide (NaN_3_) and the ATP synthase with oligomycin (Bogucka and Wojtczak, 1966; Chappell and Greville, 1961). We observed that *pfk-1.1* mutants exhibited hypersensitivity to sodium azide and oligomycin, similar to their acute response to lowered oxygen (Figure 1J). Our findings indicate that disruption of oxidative phosphorylation, either pharmacologically or by hypoxia, uncovers a role for glycolysis in the maintenance of synaptic vesicle clusters.

To better understand the role of glycolysis in the synaptic vesicle cycle, we pharmacologically inhibited glycolysis with 2-deoxy-D-glucose (2-DG) (Woodward and Hudson, 1954). We observed that animals exposed to 2-DG phenocopied *pfk-1.1* mutant animals, exhibiting hypersensitivity to oligomycin, sodium azide, and hypoxia in the maintenance of synaptic vesicle clusters (Figure S3K). Moreover, addition of 2-DG to *pfk-1.1(ola72)* animals did not enhance the synaptic vesicle clustering defects of the *pfk-1.1* mutants (Figure S3K). Together our findings indicate that *in vivo*, under conditions of energy stress in which the activity of the oxidative phosphorylation pathway is decreased, the glycolytic pathway is required in *C. elegans* neurons for the maintenance of synaptic vesicle clusters.

### Endocytosis is disrupted in *pfk-1.1* mutants during hypoxia

Synaptic vesicle endocytosis is vulnerable to ATP levels, and pharmacological or genetic inhibition of glycolysis dramatically and specifically reduces endocytosis at presynaptic sites (Rangaraju et al., 2014; Wang et al., 2004). Mutants lacking the endocytic proteins display a diffuse distribution of synaptic vesicle proteins throughout the neurite even under normoxic conditions (Schuske et al., 2003) (Figures 3A-3F). This phenotype resembles that seen for glycolytic mutants under conditions of energetic stress (Figures 2C-2I). These observations suggest that the synaptic vesicle defect observed in *pfk-1.1* mutant animals could result from inhibition of endocytosis due to decreased rates of glycolysis.

**Figure 3.**
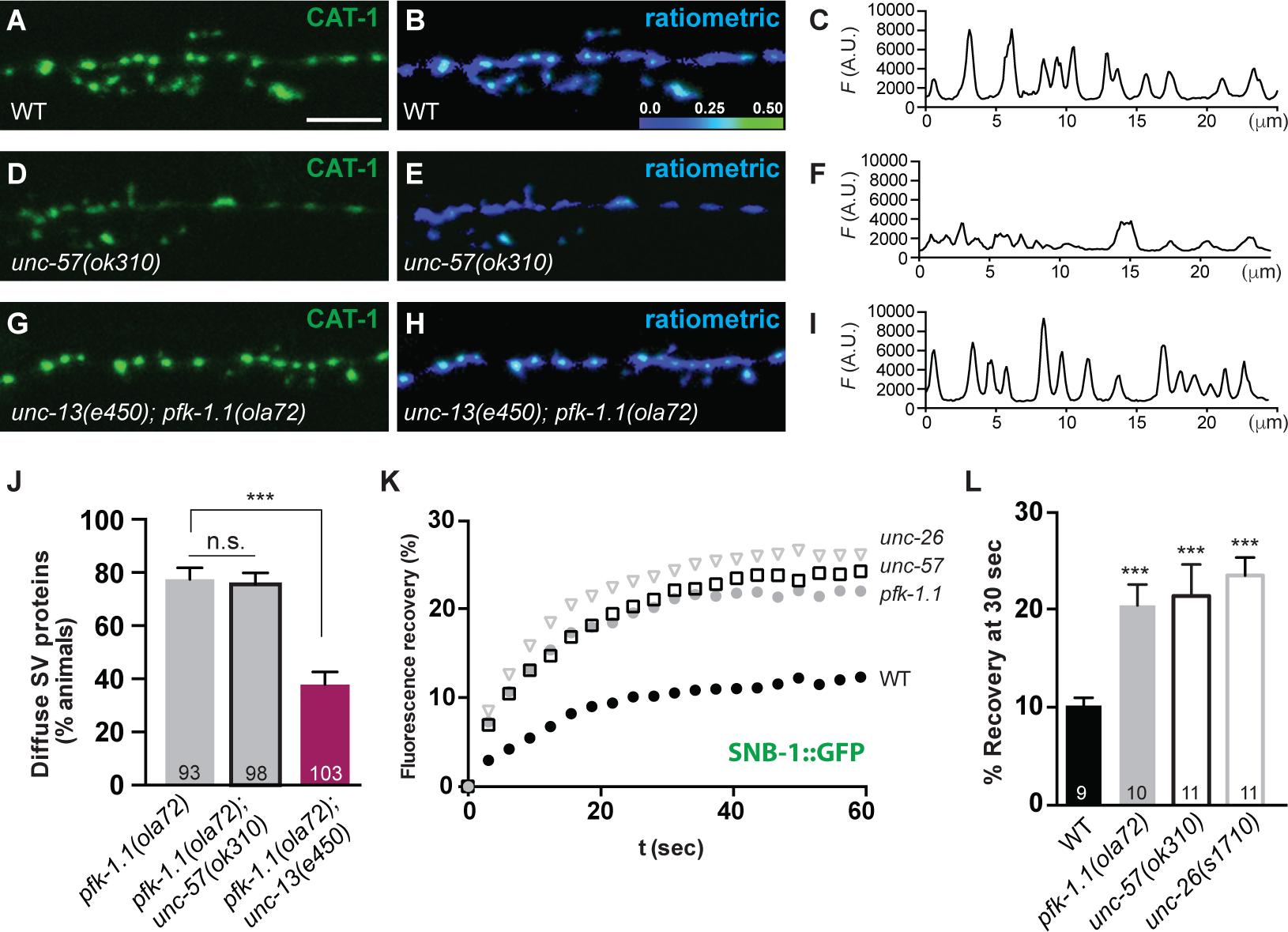
Glycolysis is required to maintain the synaptic vesicle cycle. (A-I) Synaptic vesicle marker CAT-1::GFP in NSM serotonergic neurons for wild-type after 10 minutes under hypoxic conditions (A), endocytosis mutant *unc-57(ok310)/Endophilin A* at normoxic condition (D), and *unc-13(e450);pfk-1.1(ola72)* double mutants after 10 minutes under hypoxic conditions (G). Ratiometric image of NSM representing the relative enrichment of CAT-1::GFP signal over cytosolic mCherry signal in the respective genotypes (B, E, H) and the respective line scan pixel fluorescence values of the synaptic vesicle marker CAT-1::GFP (C, F, I). (J) Percentage of animals displaying a diffuse distribution of synaptic vesicle proteins in the NSM neuron after 10 minutes of hypoxia in *pfk-1.1(ola72)* (gray bar), *unc-57(ok310); pfk-1.1(ola72)* (gray bar with black outline) and *unc-13(e450);pfk-1.1(ola72)* (red bar) double mutants. Number of animals scored is indicated at the bottom of each column. *pfk-1.1(ola72)* is the same as shown in Figure 1J. (K-L) Percentage of fluorescence recovery after photobleaching (FRAP) at synaptic varicosities of GABA neurons in *unc-26(s1710)* (triangles), *unc-57(ok310)* (squares), *pfk-1.1(ola72)* (gray circles), and wild-type (black circles) animals over time (K); and at 30 seconds post photobleaching (L). Number of animals tested is indicated at the bottom of each column. Scale bar in (A) represents 5μm. Error bars denote s.e.m. *, p < 0.05. **, p < 0.01. ***, p < 0.001.

To examine if *pfk-1.1* mutants disrupt synaptic vesicle endocytosis, we first generated double mutants of both *pfk-1.1* and endophilin/*unc-57*. Mutations in endophilin did not enhance the hypoxia-induced *pfk-1.1* phenotype (Figure 3J). Second, to examine if the vesicle proteins are trapped on the plasma membrane in the *pfk-1.1* mutants, we analyzed the mobility of vesicle protein synaptobrevin using Fluorescence Recovery After Photobleaching (FRAP). Proteins associated with the plasma membrane exhibit greater mobility than proteins associated with synaptic vesicles (Bai et al., 2010; Kraszewski et al., 1996). In the wild-type, 10% of synaptobrevin was recovered in the photobleached area by 30 seconds, whereas approximately 20% was recovered in *pfk-1.1* mutants. Endocytic mutants endophilin/*unc-57* and synaptojanin/*unc-26* exhibited similarly higher percent recoveries as *pfk-1.1* mutants (Figures 3K and 3L). Third, to test if the accumulation of vesicle proteins in the plasma membrane requires exocytosis, we blocked exocytosis by using mutants for the synaptic vesicle docking protein UNC-13. Decreased synaptic vesicle fusion in the double mutant (*pfk-1.1(ola72); unc-13(e450)*) partially suppressed the hypoxia-induced *pfk-1.1* accumulation of synaptic vesicle proteins on the surface of the axon (Figures 3G-3J). The observed partial suppression may be due to the hypomorphic nature of the *unc-13* allele, or to potential contributions to the phenotype from other, *unc-13* independent pathways. The phenotype in *pfk-1.1; unc-13* double mutants, however, indicates that a defect in the synaptic vesicle cycle significantly contributes to the *pfk-1.1* mutant phenotype. Our results are consistent with previous studies (Rangaraju et al., 2014; Wang et al., 2004) and suggest that under conditions of energy depletion, the glycolytic pathway becomes critical in maintaining the energy supplies necessary for sustaining endocytosis and the synaptic vesicle cycle.

### Glycolytic enzymes localize to presynaptic sites during energy stress

Where is PFK-1.1 localized to meet local energy demands at synapses? We examined the subcellular localization of PFK-1.1 in neurons. We observed that under normoxic conditions, PFK-1.1 is localized in a punctate pattern at some of the cell somas (Figure S4A) and largely diffuse throughout the neurites (Figures 4A and 4C). However, upon exposure to hypoxia, PFK-1.1 became clustered in neurites (Figures 4A-4E; Movie S2), preferentially localizing to synaptic rich regions (Figure 4F) and within 0.2 um from vesicle release sites (Figures 4G and 4H).

**Figure 4.**
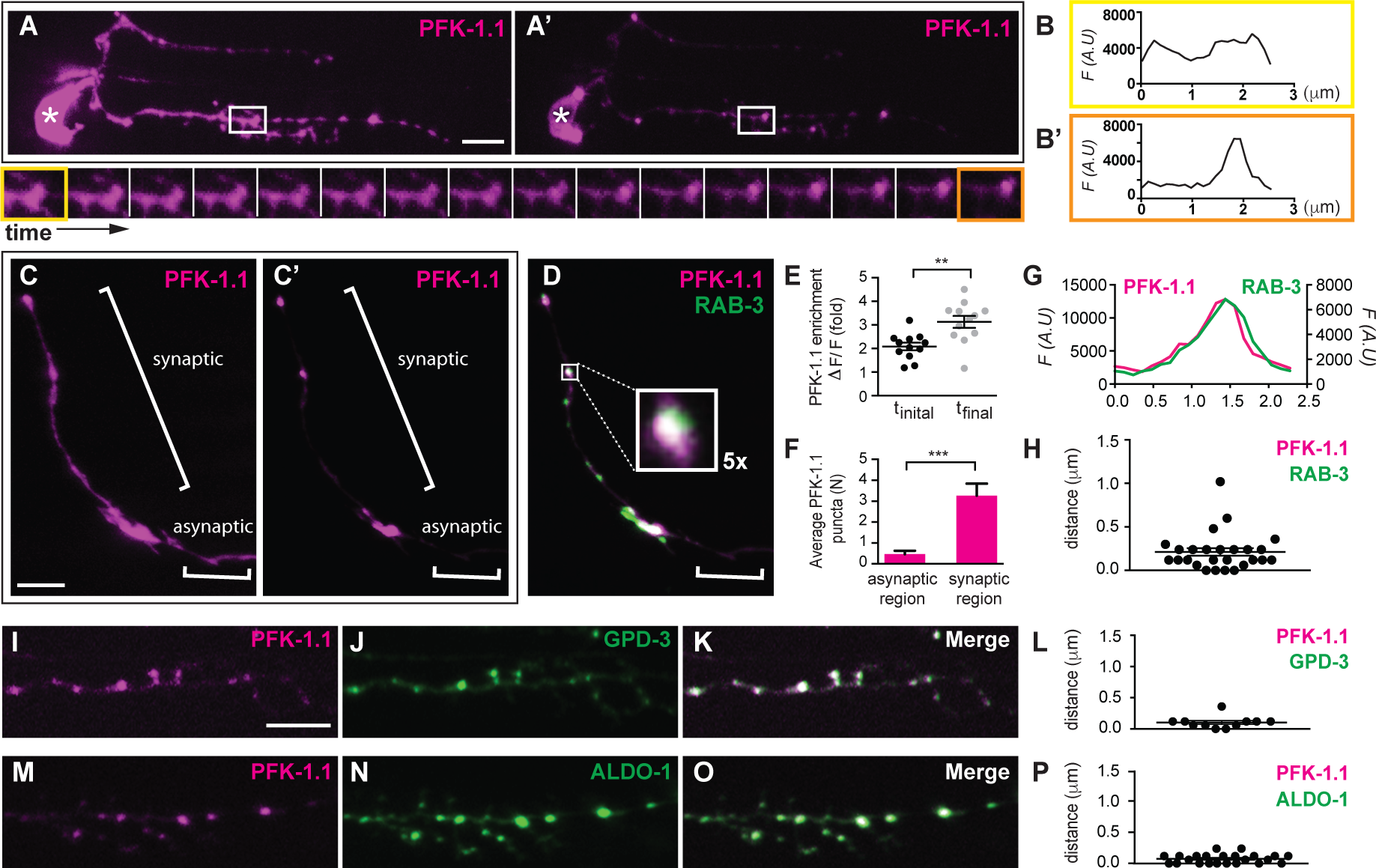
Glycolytic proteins dynamically clusters near presynaptic release sites. (A-A’) PFK-1.1::eGFP (pseudocolored magenta) localization in NSM after approximately 10 minutes (A) and 30 minutes (A’) of exposure to hypoxic conditions (see *Experimental Procedures*). Clustering dynamics of PFK-1.1 vary between individual animals, but clustering of PFK-1.1 is observed for most animals within 10 minutes of being exposed to hypoxic conditions. Here we filmed a neuron with slow clustering dynamics to capture the time-lapse image. Cell body of NSM neuron is indicated by asterisk. The time-lapse images of the inset in (A) and (A’) (taken at 1 minute intervals) are displayed below, starting at approximately minute 10 and extending to minute 30. See Movie S2 for time-lapse movie. (B-B’) Pixel fluorescence values for the first inset in the montage (B) (yellow trim in bottom row) and the last inset (B’) (orange trim in bottom row). (C-E) PFK-1.1::eGFP (pseudocolored magenta) and mCherry::RAB-3 (pseudocolored green) simultaneously visualized in AIY interneurons. PFK-1.1 in AIY after 20 minutes (C); and 40 minutes (C’) of hypoxic conditions, and C’ visualized with synaptic vesicle marker RAB-3 (D); the brightness of (C) was increased more than in (C’) in order to allow for visibility of the diffuse localization of PFK-1.1. Note that PFK-1.1 becomes increasingly clustered to presynaptic sites (denoted with brackets as “synaptic region”), and is excluded from asynaptic regions (Colon-Ramos et al., 2007; White et al., 1986). Inset in (D) corresponds to zoomed-in (5x) image of PFK-1.1 cluster localizing near a pool of synaptic vesicles. Enrichment of PFK-1.1 clustering in the AIY neurite over time in hypoxic treatment was quantified (N=12) (See *Experimental Procedures*) (E). t_initial_ is from images of less than 20 minutes of hypoxic treatment and t_final_ is from images between 20 to 40 minutes of hypoxic treatment, and each circle represents and individual animal in the longitudinal study. (F) Quantification of the number of PFK-1.1 clusters in the synaptic region and the asynapticregions of AIY (N=15). Note that PFK-1.1 clustering preferentially occurs at the synapticregions. (G) Pixel fluorescence values of PFK-1.1 and RAB-3 along the neurite for boxed region in D tocompare the colocalized distribution of PFK-1.1 relative to synaptic vesicle proteins. (H) Quantification of the distance between the max pixel fluorescence values of PFK-1.1 and the max pixel fluorescence values of RAB-3 in AIY neurons (N=26). (I-P) PFK-1.1::mCherry (pseudocolored magenta) (I, M); and glycolytic proteins GPD-3::eGFP (J); or ALDO-1::eGFP (N) were simultaneously visualized in NSM neurons. GDP-3::eGFP and ALDO-1::eGFP were distributed throughout the cytosol under normoxic conditions, similar to PFK-1.1 (data not shown). After 10 minutes of hypoxic conditions, GDP-3::eGFP and ALDO-1::eGFP were observed to cluster (J, N); and colocalize with PFK-1.1::mCherry (K, O). Quantification of the distance between the max pixel fluorescence values of PFK-1.1 and GPD-3 (L) (N=11) and PFK-1.1 and ALDO-1 (P) (N=23). Scale bar in all panels represents 5μm. Error bars denote s.e.m. *, p < 0.05. **, p < 0.01. ***, p < 0.001.

Glycolytic proteins have been hypothesized to interact and form functional super-complexes, termed glycolytic metabolons (or glycolons), which sustain the accelerated rates of glycolysis (Clarke and Masters, 1975; Kurganov et al., 1985; Ureta, 1985). The concept of the glycolytic metabolon derives from three lines of evidence generated primarily from work in fixed tissues or biochemical assays: 1) enzymatic and modeling studies suggesting that glycolytic enzymes need to be near one another to sustain the observed enzymatic rates (Boiteux and Hess, 1981; Clarke and Masters, 1975; Kurganov et al., 1985; Zhou et al., 2005); 2) immunohistological studies demonstrating colocalization of glycolytic proteins to subcellular structures in specific cell types (Campanella et al., 2008; Raikar et al., 2006; Saez and Slebe, 2000; Sullivan et al., 2003); and 3) biochemical studies indicating increased interactions between glycolytic enzymes under conditions of energy stress (Bronstein and Knull, 1981; Knull et al., 1980; Masters, 1984; Mowbray and Moses, 1976; Puchulu-Campanella et al., 2013). Less is known about the existence, *in vivo*, of this complex or its physiological importance (Menard et al., 2014). To examine if other glycolytic enzymes colocalize with PFK-1.1 *in vivo*, we visualized the subcellular localization of ALDO-1 and GPD-3. ALDO-1 and GPD-3, like PFK-1.1, were diffusely distributed throughout the cytosol under normoxic conditions (data not shown). Upon exposure to hypoxia, both enzymes clustered near synapses (Figures 4J and 4N). The GPD-3 and ALDO-1 clusters colocalized with the PFK-1.1 clusters (Figures 4I-4P), suggesting that these enzymes dynamically form a complex near presynaptic sites in response to demands for ATP.

Aerobic glycolysis normally occurs in the brain, and increases locally upon conditions of increased neuronal activity (Vaishnavi et al., 2010). To test if the glycolytic metabolon is necessary at synapses in response to high levels of neuronal activity we stimulated neurons using a pharmacological approach. *C. elegans* GABA neurons can be stimulated through the bath application of levamisole, an agonist of acetylcholine receptors (such as *lev-8*) present in the GABA neurons (Towers et al., 2005). By applying levamisole under normoxic conditions, we examined whether *pfk-1.1* was required to support vesicle cycling during periods of strong neuronal stimulation. We observed that *pfk-1.1* mutant animals were sensitized to levamisole treatment compared to wild-type animals (Figures 5A-5C). This sensitization is specific to the GABA neurons that express levamisole-sensitive acetylcholine receptors (data not shown). These findings suggest that even in aerobic conditions, glycolytic proteins are required to maintain the synaptic vesicle cycle during periods of high neuronal activity, which are known to result in increased energy demands at synapses (Rangaraju et al., 2014). Together, our findings suggest that glycolysis is critical for the maintenance of the synaptic vesicle cycle under conditions that either reduce energy supplies (such as hypoxia), or increase energy demands (such as neuronal stimulation).

**Figure 5.**
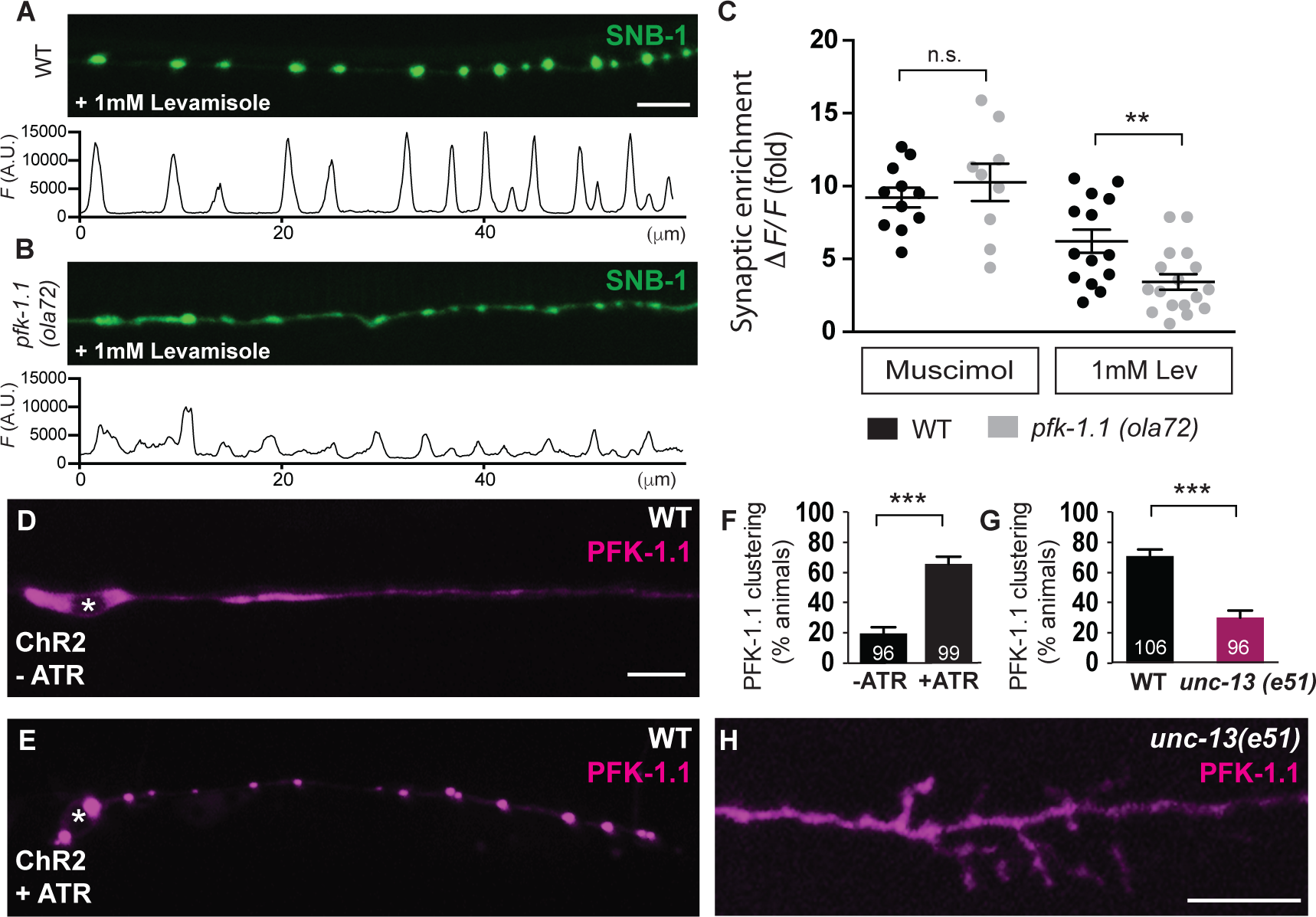
Neuronal stimulation induces PFK-1.1 clustering. (A-B) Synaptic vesicle protein SNB-1::GFP in GABA neurons exposed to levamisole in wild-type (A); or *pfk-1.1(ola72)* mutant animals (B). GABA neurons express acetylcholine receptors that are stimulated by the cholinergic agonist levamisole (Towers et al., 2005). Pixel fluorescence values of the synaptic vesicle proteins along the neurites are shown under each (A) and (B). (C) Quantification of the synaptic enrichment of the synaptic vesicle proteins SNB-1, (*ΔF/F*) in the GABA neurons for wild-type (black circles) and *pfk-1.1(ola72)* (gray circles) under muscimol or levamisole treatment. The circles in the graph represent individual animals. GABA agonist muscimol does not stimulate GABA neurons (Schuske et al., 2004) while the cholinergic agonist levamisole stimulates GABA neurons (Towers et al., 2005). Under levamisole treatment, *pfk-1.1(ola72)* mutant animals (gray circles) has lower synaptic enrichment (more diffuse distribution of the synaptic vesicle proteins) compared to the wild-type (black circles). The response of GABA neurons to levamisole is dependent on the acetylcholine receptor LEV-8 (Towers et al., 2005), and was suppressed in *lev-8(ok1519)* mutant animals (data not shown). (D-E) Localization of PFK-1.1 (pseudocolored magenta) in wild-type animals expressing channelrhodopsin cell-specifically in GABA neurons and stimulated with blue light in the absence (D); or presence (E) of rhodopsin co-factor all-trans-retinal (ATR). Cell body is indicated by asterisk. (F) Percentage of animals displaying PFK-1.1 clusters in GABA neurites expressing channelrhodopsin and stimulated with blue light in the absence or presence of ATR. Number of animals tested is indicated at the bottom of each column. (G) Percentage of animals displaying PFK-1.1 clusters in NSM neurites under hypoxic conditions in wild-type (black bar) and *unc-13(e51)* (red bar) mutant backgrounds. Number of animals tested is indicated at the bottom of each column. (H) Localization of PFK-1.1 in NSM neurons of *unc-13(e51)* animals exposed to 10 minutes of hypoxic conditions, quantified in (G). Scale bar represents 5μm. Error bars denote s.e.m. *, p < 0.05. **, p < 0.01. ***, p < 0.001 between indicated groups.

Does PFK-1.1 cluster at synapses upon increased neuronal stimulation? Optogenetic stimulation of GABA neurons expressing channelrhodopsin caused clustering of PFK-1.1 in normoxic conditions (Figures 5D-5F). Therefore PFK-1.1 can dynamically localize near presynaptic sites upon neuronal stimulation. To test whether this relocalization is dependent on metabolic needs at the synapse, we examined, in hypoxic conditions, PFK-1.1 localization in *unc-13* mutants, in which exocytosis and synaptic activity are greatly reduced (Richmond et al., 1999). We observed that the subcellular localization of PFK-1.1 to presynaptic sites is reduced in the neurites (but not the somas) of *unc-13(e51)* mutants (Figures 5G, 5H, and S4A). In normoxic conditions, optogenetic stimulation of GABA neurons did not promote PFK-1.1 clustering in *unc-13* mutants, indicating that synaptic PFK-1.1 clustering does not simply depend on depolarization (Figure S4B). These results suggest that PFK-1.1 clustering to presynaptic sites relies on the synaptic vesicle cycle and energy demands at synaptic sites.

### Presynaptic localization of PFK-1.1 depends on synaptic scaffolding proteins

How do glycolytic proteins localize to presynaptic sites? Glycolytic proteins co-purify with synaptic vesicles and are required for their transport (Burre and Volknandt, 2007; Ikemoto et al., 2003; Ishida et al., 2009; Zala et al., 2013). Glycolytic proteins are also known to associate with the mitochondria, which are actively transported to synaptic sites (Giege et al., 2003). Therefore we next examined if the localization of glycolytic proteins to presynaptic sites depended on synaptic vesicle transport or on mitochondrial transport to synapses. We observed that in *unc-104/kinesin 3/kif1A* mutant animals, synaptic vesicles fail to cluster at presynaptic sites, as previously reported (Hall and Hedgecock, 1991; Nelson and Colón-Ramos, 2013). However, PFK-1.1 clustering under hypoxic conditions was not affected in *unc-104(e1265)* mutant animals (Figures 6A-6D, and 6O). Our data indicate that the clustering of PFK-1.1 at presynaptic sites is not dependent on vesicle transport to synapses.

**Figure 6.**
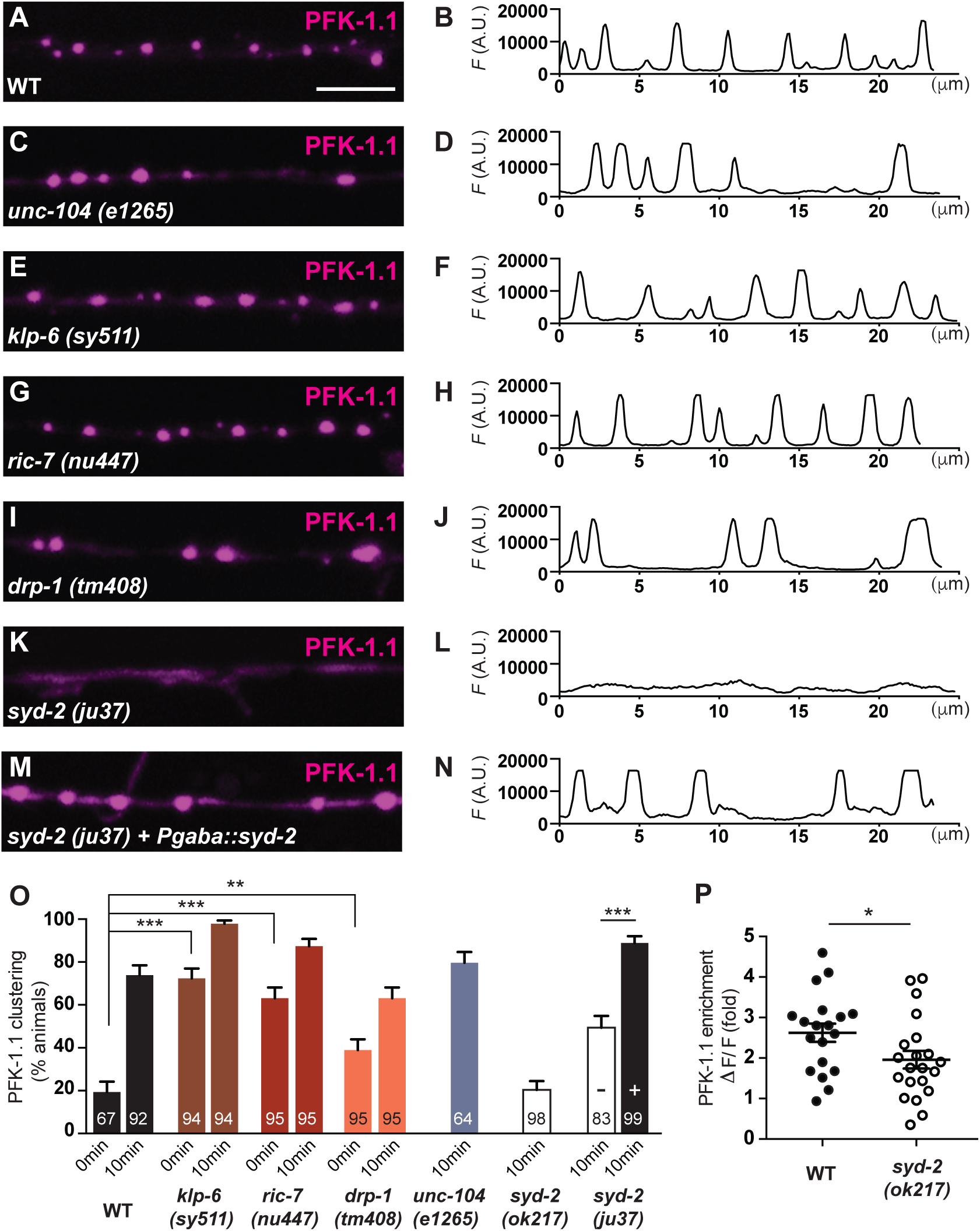
Presynaptic localization of PFK-1.1 depends of synaptic scaffolding proteins. (A-N) Localization of PFK-1.1 in GABA ventral neurites (pseudocolored magenta) and its respective pixel fluorescence values along the neurite after 10 minutes of hypoxia in wild-type (A, B); synaptic vesicle kinesin mutant *unc-104(e1265)* (C, D); mitochondria localization and distribution mutants *klp-6(sy511)* (E, F), *ric-7(nu447)* (G, H), and *dpr-1(tm408)* (I, J); synaptic scaffolding mutant *syd-2(ju37)* (K, L); and *syd-2(ju37)* mutant animals expressing a wild-type rescuing array of *syd-2* cell-specifically in GABA neurons (M, N). (O) Percentage of animals displaying PFK-1.1 clusters in GABA neurites in varying genotypes after 0 or 10 minutes of exposure to hypoxic conditions, as indicated. For *syd-2(ju37)*, PFK-1.1 clustering was examined in the absence (–) or in the presence (+) of a wild-type rescuing array of *syd-2* cell-specifically in GABA neurons. Number of animals tested is indicated at the bottom of each column. (P) Quantification of the synaptic enrichment of PFK-1.1, (*ΔF/F*) in the NSM neurons for wild-type (black circles) and *syd-2(ok217)* (white circles) under hypoxic conditions. The circles in the graph represent individual animals. Scale bar represents 5μm. Error bars denote s.e.m. *, p < 0.05. **, p < 0.01. ***, p < 0.001 between indicated groups.

To determine if the localization of PFK-1.1 to presynaptic sites depends on the mitochondria, we simultaneously visualized the subcellular localization of PFK-1.1 and mitochondrial outer-membrane protein TOM-20 in mutant backgrounds that affect mitochondria localization in neurites. We observed that in the mutant backgrounds of kinesin *klp-6(sy511)*, *ric-7(nu447)* and mitochondrial fission *drp-1(tm408)* there was a reduction (or altered distribution) of mitochondria in the neurites, as previously reported (data not shown) (Labrousse et al., 1999; Rawson et al., 2014; Tanaka et al., 2011). While the mitochondria localization was affected in these animals, we did not detect a reduction in the number of animals displaying PFK-1.1 clusters in the neurites (Figures 6E-6J, and 6O). Instead, we observed that *klp-6(sy511)*, *ric-7(nu447)*, and *drp-1(tm408)* mutants enhanced PFK-1.1 clustering in neurites even before animals were exposed to hypoxic conditions (Figure 6O). Our observations that PFK-1.1 clustering is observed in these mitochondria mutants even before animals are exposed to hypoxic conditions are consistent, and extend our findings on the role of glycolysis under conditions of energy stress. Importantly, our findings indicate that glycolytic protein localization to presynaptic sites is not dependent on the mitochondria or *unc-104/kinesin 3/kif1A* synaptic vesicle transport.

SYD-2 is a synaptic scaffolding protein required for synaptic release sites formation and maintenance (Zhen and Jin, 1999). To examine if the integrity of the presynaptic sites was necessary for the localization of glycolytic clusters in the neurite, we visualized PFK-1.1 in *syd-2* mutant animals. We observed that in the *syd-2(ok217)* null allele (Wagner et al., 2009) and the *syd-2(ju37)* loss of function allele (Zhen and Jin, 1999), PFK-1.1 clustering at the neurites was significantly suppressed (Figures 6K, 6L, 6O, and 6P). Cell-specific expression of the wild-type SYD-2 in GABA neurons rescued the PFK-1.1 clustering in GABA neurons of *syd-2 (ju37)* mutants (Figures 6M-6O), demonstrating the cell autonomous role of this synaptic scaffolding protein in PFK-1.1 localization. Similar results were obtained for presynaptic protein SYD-1 (data not shown) (Hallam et al., 2002). Our findings demonstrate that presynaptic scaffolding proteins are cell autonomously required in neurons for the clustering of PFK-1, and underscore a functional link between the presynaptic release sites and the clustering of glycolytic proteins.

### Localization of PFK-1.1 to synaptic sites is important for maintaining the synaptic vesicle cycle during energy stress

To test the importance of the subcellular localization of PFK-1.1 in sustaining energy levels at synapses, we identified a dominant negative version of PFK-1.1 (PFK-1.1G532E). PFK-1.1G532E contains a lesion in the regulatory domain of PFK-1.1, fails to form clusters (Figures S4C-S4E), and fails to rescue the *pfk-1.1* mutant phenotype (data not shown). We observed that expression of PFK-1.1G532E in wild-type animals altered wild-type PFK-1.1 localization to presynaptic sites, suggesting that PFK-1.1G532E acts as a dominant negative, affecting the subcellular localization of wild-type PFK-1.1 (Figures 7A-7D). In addition, these wild-type animals expressing PFK-1.1G532E display a dominant negative synaptic vesicle clustering phenotype similar to that observed for *pfk-1.1* loss-of-function mutant animals (Figures 7E-7I). Because expression of PFK-1.1G532E affects both wild-type PFK-1.1 localization and function, we hypothesize that disruption of the localization of the wild-type PFK-1.1 affects its capacity to sustain energy levels at the synapse, and the synaptic vesicle cycle. Consistent with these findings, we also observed that when we alter the localization of a wild-type version of PFK-1.1 with a nuclear localization sequence, the wild-type PFK-1.1 is incapable of rescuing the synaptic vesicle phenotype of *pfk-1.1* mutants (Figures S4F-S4H). Our findings are consistent with studies in *Drosophila* fly muscles, which demonstrated that even when the full complement of glycolytic enzymes is present in the muscle, disruption of glycolytic protein colocalization to sarcomeres results in inability to fly (Wojtas et al., 1997). Together our findings suggest that the presence of the enzymes in the neurons is not sufficient to power synaptic function. Instead, subcellular localization of these glycolytic enzymes near synapses is necessary to power synaptic function.

**Figure 7.**
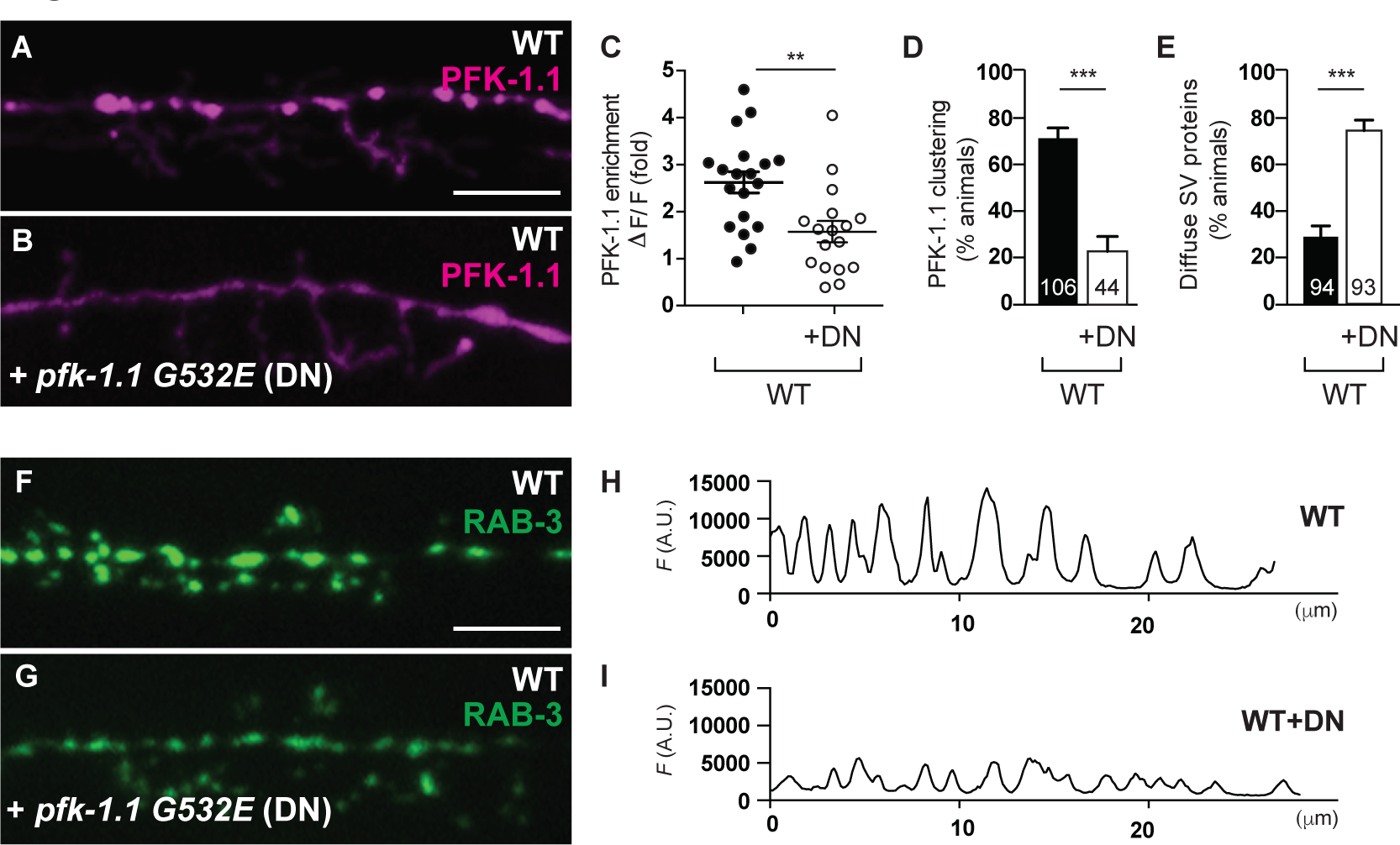
Localization of PFK-1.1 is necessary for maintaining vesicle clusters under energy stress. (A-B) Wild-type NSM neuron in the absence or presence of PFK-1.1(G532E) (dominant negative, DN). In the absence of PFK-1.1(G532E), PFK-1.1::eGFP (pseudocolored magenta) localizes, and is enriched at presynaptic sites under energy stress (A) as also shown in Figures 4 and 5E. However, expression of PFK-1.1(G532E) prevents the enrichment of wild-type PFK-1.1 at presynaptic sites (B). PFK-1.1(G532E) also fails to localize to presynaptic sites (Figures S4C-4E). (C) Quantification of the enrichment of PFK-1.1::eGFP WT (black circles, N=19), and PFK-1.1:eGFP WT co-expressed with the dominant negative PFK-1.1(G532E) (+DN and white circles, N=17). Wild-type control is the same as shown in Figure 6P. (D) Percentage of animals displaying PFK-1.1 clusters or diffuse distribution of synaptic vesicle proteins (E) in wild-type (WT) NSM neurons under hypoxic conditions when expressed without (black bar) or with PFK-1.1(G532E) (+DN and white bar). Number of animals scored is indicated at the bottom of each column. Wild-type control is the same as shown in Figure 5G. (F-I) Synaptic vesicle proteins RAB-3 (pseudocolored green) in NSM neurons and its respective pixel fluorescence along the NSM neurite in wild-type animals (F, H) and in wild-type animals expressing the dominant negative (DN) PFK-1.1 (G532) (G, I) under hypoxic conditions. The respective penetrance is quantified in (E). Scale bar represents 5μm. Error bars denote s.e.m. *, p < 0.05. **, p < 0.01. ***, p < 0.001 between indicated groups.

### PFK-1.1 is required to sustain synaptic activity and locomotion during energy stress

Is PFK-1.1 required to sustain synaptic activity? We examined the requirement of *pfk-1.1* in synaptic recovery following fatigue. Animals expressing the channelrhodopsin variant ChIEF were dissected to expose neuromuscular synapses of the body muscles (Richmond et al., 1999; Richmond and Jorgensen, 1999). Postsynaptic muscles were voltage-clamped and high-frequency pulses of blue light were delivered to stimulate synapses for 30 seconds at 10Hz. This stimulation caused rapid fatigue of acetylcholine neuromuscular junctions (Liu et al., 2009), presumably by the depletion of vesicle pools. We applied test-pulses at increasing time increments after fatigue and assayed the recovery of evoked currents (Figures 8A and 8B). We observed that the absolute value of the first evoked response in the *pfk-1.1* mutant is similar to the wild-type, demonstrating that the synapses of both wild-type and *pfk-1.1* mutants are healthy and normal in response to an evoked stimulus (Figure 8C). Moreover, under normoxic conditions, both wild-type and *pfk-1.1* mutant synapses recovered with a time constant of approximately 9 seconds, achieving full recovery (Figure 8D). However, with two and three repetitions of the 30 second stimulation protocol, *pfk-1.1* mutants exhibited reduced recovery (72% and 40% of wild-type levels, respectively) (Figures 8E and 8F). Next we included oligomycin to block oxidative phosphorylation. The rate of recovery at wild-type synapses was slightly decreased but not significantly different from control conditions (time constant of 12 seconds). Furthermore, the drug did not affect the maximum recovery (Figure 8G). By contrast, *pfk-1.1* mutants recovered more slowly (time constant >35 seconds) and failed to reach full recovery (60% of wild-type plateau amplitude) (Figure 8G). Therefore oxidative phosphorylation or glycolysis alone support ATP demands under moderate stimulation. However, glycolysis is required for synaptic function under persistent stimulus conditions, or in the absence of oxidative phosphorylation.

**Figure 8.**
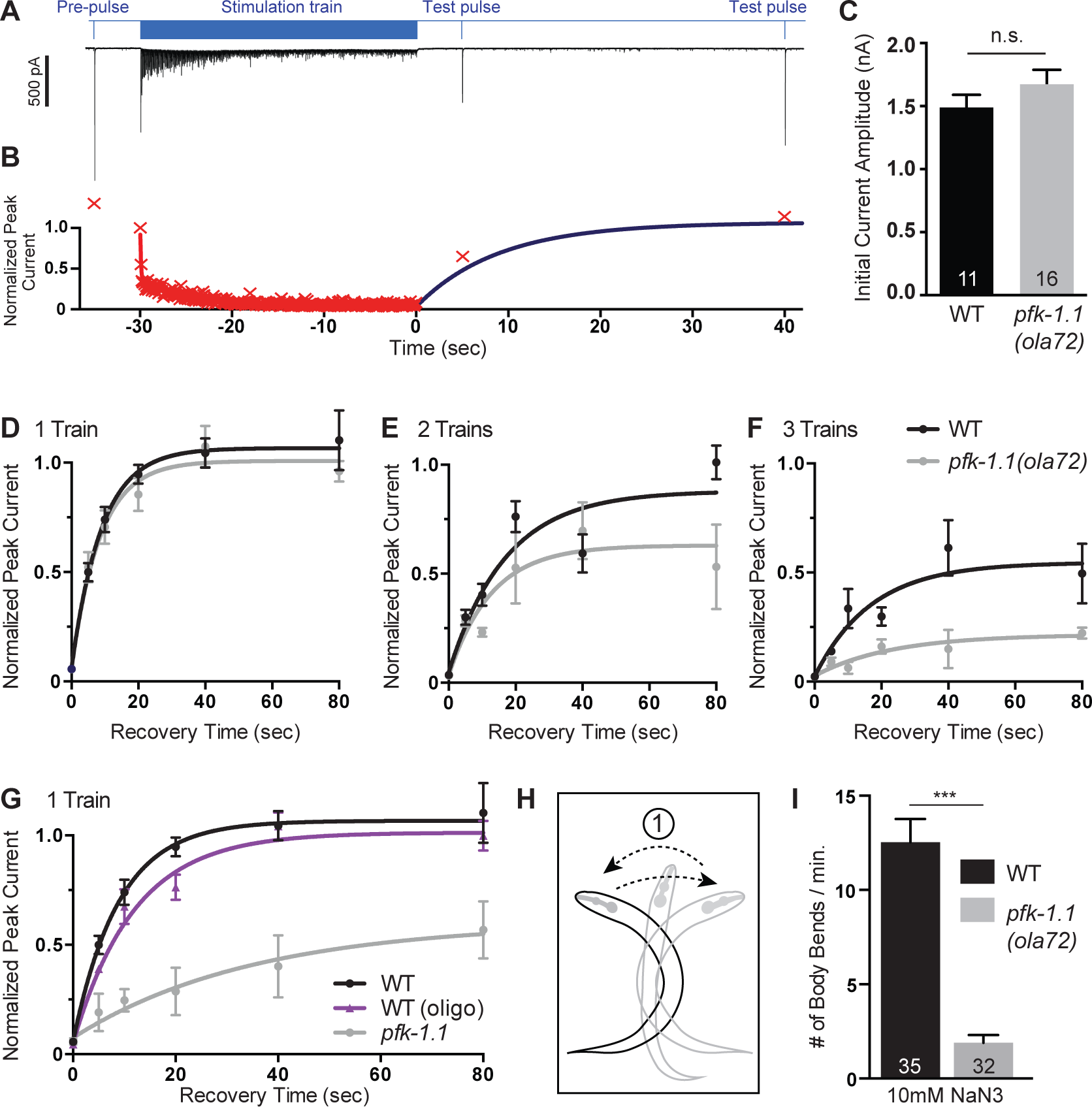
PFK-1.1 is required for synaptic recovery following fatigue and for locomotion. (A) Representative electrophysiology experiment illustrating synaptic fatigue and recovery at *C. elegans* neuromuscular junctions. Experiments were conducted in strains expressing channelrhodopsin in acetylcholine neurons *oxSi91* [P*unc-17::ChIEF*] that lack GABA inputs to the muscle (*unc-49(e407)*). Animals were dissected and body muscles were voltage-clamped. The blue line indicates blue-light illumination and the black trace below is an example trace plotting postsynaptic current. A single pre-pulse that eliminates further inactivation was applied, and followed by a 5-second recovery period and 30-second stimulation train (10Hz). To assay recovery, two test pulses (>15 seconds apart) were delivered at fixed times after the stimulation train. (B) Normalized plot of representative evoked amplitudes. Peak current was measured for each evoked response (red Xs) and normalized to the amplitude of the first pulse of the stimulus train. The evoked amplitudes from this experiment are plotted vs. time. The fitted curve for recovery in wild-type animals (dark blue curve) is overlaid on the data points. In subsequent panels only the recovery data is presented. (C) Mean peak current for the first stimulus in the train for wild-type (black bar) and *pfk-1.1(ola72)* mutants (gray bar). Number of animals scored is indicated at the bottom of each column. (D) Synaptic recovery of *pfk-1.1(ola72)* mutants is equivalent to wild-type recovery following one stimulus train. Wild-type and *pfk-1.1(ola72)* mutant strains reached full recovery (plateau amplitude +/-SEM: WT=1.1 +/-0.037; *pfk-1.1* =1.0 +/-0.046) with time constants of 8.9 and 8.6 seconds respectively. (N=19 animals for the wild-type; N=14 animals for *pfk-1.1*) (E-F) Synaptic recovery is reduced in *pfk-1.1(ola72)* mutants following two (D) and three (E) 30 second stimulus trains. When the stimulus protocol was delivered twice, *pfk-1.1(ola72)* recovery decreased to 72% of wild-type recovery (plateau amplitude: WT=0.88 +/-0.071; *pfk-1.1* =0.63 +/-0.086) (N=12 animals for wild-type; N=9 animals for *pfk-1.1*)(D). With three stimulus trains, the plateau amplitude was reduced to 40% of wild-type levels (WT=0.55 +/-0.071; *pfk-1.1* =0.22 +/-0.043)(**e**). N=10 animals for wild-type; N=8 animals for *pfk-1.1*. (G) Synaptic recovery is significantly reduced in *pfk-1.1(ola72)* mutants in the presence of oligomycin following one stimulus train. Oligomycin (1uM oligomycin for 5 min) did not significantly affect synaptic recovery in wild-type animals (time constant: oligo=11.9 sec vs. control=8.9 sec; plateau amplitude: oligo=1.0 +/-0.047 vs. control=1.1 +/-0.037). *pfk-1.1* mutant animals recovered 3-times slower than wild-type animals (time constant: *pfk-1.1* >35 sec vs. WT = 12 sec) and never reached full recovery (plateau amplitude: 0.60 +/- 0.17), and amplitudes were significantly different from the wild-type (p<0.05 Mann-Whitney *U* test; for t=10, 20, and 40 seconds; n=3-7 animals per time point for the wild-type; n=4-7 animals per time point for *pfk-1.1*; N=12 animals for wild-type; N=16 animals for *pfk-1.1*). (H) Cartoon illustration of one full body bend in *C. elegans* as indicated by the arrows. (I) Number of body bends measured for 1 minute after a four-minute bath in a 10mM sodium azide (NaN_3_). *pfk-1.1(ola72)* mutants show a significant reduction in the number of body bends as compared to wild-type animals. Number of animals scored is indicated at the bottom of each column. M9 buffer alone produced no difference in the number of body bends between wild-type and *pfk-1.1(ola72)* mutant animals (not shown). Error bars denote s.e.m. *, p < 0.05. **, p < 0.01. ***, p < 0.001 between indicated groups.

To determine if glycolysis is required for normal behavior, we tested whether *pfk-1.1* mutant animals exhibited fatigue during swimming. Worms thrash rapidly in liquid, so we scored the number of body bends in solution per minute (Figure 8H). We observed no differences in the number of body bends between wild-type and *pkf-1.1* mutants under ambient conditions (data not shown). To inhibit oxidative phosphorylation, animals were placed in solution containing 10mM sodium azide (NaN_3_) or under hypoxic conditions. Consistent with synaptic impairments in *pfk-1.1* mutants, we observed a significant reduction in the number of body bends in *pfk-1.1* mutants compared to wild-type animals (Figure 8I and data not shown). Together, these data suggest that disruption of the local formation of the glycolytic metabolon results in synaptic impairment and behavioral deficits *in vivo*.

## Discussion

While many cellular processes rely on ATP, processes are differentially vulnerable to decreases in ATP production, depending on their K_m_ values (Rangaraju et al., 2014). A recent study demonstrated that synaptic vesicle endocytosis is a particularly ATP-sensitive process and more vulnerable to metabolic perturbations than exocytosis (Rangaraju et al., 2014). Genetic studies in *Drosophila* also identified a role for the glycolytic enzyme phosphoglycerate kinase in regulating synaptic transmission, and demonstrated that the physiological phenotypes of phosphoglycerate kinase mutants were related to problems in synaptic vesicle endocytosis due to reduced ATP levels (Wang et al., 2004). Our findings in *C. elegans* neurons now demonstrate that the glycolytic machinery is required *in vivo* for the endocytosis of synaptic vesicle proteins under conditions of energetic stress.

In most organisms, glycolytic proteins are soluble in the cytosol. An exception to this is the protozoan *Trypanosoma brucei*, which organizes its glycolytic enzymes in a membrane-bound organelle called the glycosome (Michels et al., 2006). This compartmentalization is essential for the regulation of the trypanosomatids’ metabolism and viability (Haanstra et al., 2015). Although membrane-bound glycosomes have not been observed in mammalian cells, it has been predicted for over thirty years that glycolytic proteins compartmentalize in the cytosol into a “glycolytic metabolon” to sustain the observed rates of glycolysis (Clarke and Masters, 1975; Kurganov et al., 1985; Ureta, 1985). *In vivo* evidence for the existence of this complex, and its physiological importance, has been lacking (Brooks and Storey, 1991; Menard et al., 2014).

We observe in *C. elegans* that glycolytic proteins dynamically cluster near presynaptic sites under conditions of energy stress, suggesting the *ad hoc* formation of a glycolytic metabolon. Disruption of presynaptic scaffolding proteins, or of the synaptic vesicle exocytosis protein, *unc-13*, suppresses the clustering of glycolytic proteins near presynaptic sites (but not their clustering at cell somas). Our findings are consistent with biochemical studies that demonstrated an enrichment of glycolytic enzymes and activity in synaptosomal fractions and lysed nerve endings (Knull, 1978; Knull and Fillmore, 1985; Wu et al., 1997), and extend findings demonstrating the presence of glycolytic proteins at postsynaptic sites and their role in synaptic transmission (Laschet et al., 2004). Interestingly, localization of glycolytic proteins to presynaptic sites does not depend on the synaptic vesicle transport protein UNC-104/Kinesin 3/kif1A. These finding indicate that clustering of glycolytic proteins to presynaptic sites is not dependent on active synaptic vesicle transport. Our findings also indicate that clustering of glycolytic proteins to presynaptic sites might also be powering other synaptic events besides the recycling of *unc-104*-dependent synaptic vesicles. Together our study demonstrates a link between the dynamic localization of glycolytic proteins *in vivo* and their functional requirement at presynaptic sites.

The formation of the glycolytic metabolon meets local energy demands at presynaptic sites. Disrupting the formation of the glycolytic metabolon at synapses —by forcing the localization of PFK-1.1 to the nucleus or using a dominant negative version of PFK-1.1— disrupts the capacity of the glycolytic pathway to sustain the synaptic vesicle cycle, synaptic physiology and locomotory activity of animals under energy stress. Our observations in *C. elegans* neurons are consistent with studies in *Drosophila* muscles that demonstrated that colocalization of glycolytic proteins to sarcomeres was necessary for muscular function (Wojtas et al., 1997). Together, our findings underscore the *in vivo* importance of the local, *ad hoc* formation of glycolytic complexes near presynaptic sites in meeting energy demands and sustaining synaptic function and behavior.

Glycolysis is necessary to sustain the synaptic vesicle cycle when oxidative phosphorylation is inhibited. Mitochondria, which mediate oxidative phosphorylation, are actively transported to neuronal synapses to meet energy demands. The physiological importance of mitochondria localization is perhaps best exemplified by the fact that defects in mitochondrial localization have been linked to neurodegenerative disorders (Burte et al., 2015; Lin and Sheng, 2015; Schwarz, 2013). In this study we observe that inhibition of mitochondrial function, or of mitochondrial transport, enhances local clustering of glycolytic proteins to presynaptic sites. We hypothesize that this enhanced clustering of glycolytic proteins represents a response to energy demands at synapses in conditions of mitochondrial disruption. Our findings demonstrate that glycolysis can act redundantly with oxidative phosphorylation to sustain the synaptic vesicle cycle, particularly under conditions in which mitochondrial function at the synapse is compromised.

While many of our experiments were conducted under hypoxic conditions, we observed that pharmacological stimulation of neurons under normoxic conditions also requires glycolysis to sustain the synaptic vesicle cycle. We also observed that optogenetic stimulation of neurons under normoxic conditions results in clustering of glycolytic enzymes. This clustering is likely due to synaptic function and not mere depolarization, as it is suppressed in mutants with reduced exocytosis (*unc-13* mutants). We therefore hypothesize that in physiological conditions, aerobic glycolysis would play an important role in sustaining synaptic function.

Synapses do not consume energy at a consistent rate, but rather have extended periods of low activity punctuated by periods of intense activity. How are changing energy demands at presynaptic sites dynamically met? Mitochondrial localization is an important mechanism for meeting local energy demands at the synapse, yet many presynaptic terminals, while rich in ATP, lack mitochondria (Chavan et al., 2015; Waters and Smith, 2003; Xu-Friedman et al., 2001). The capacity of the glycolytic machinery to produce ATP molecules at a faster rate than oxidative phosphorylation, and to dynamically assemble into metabolic compartments based on energy needs, might fill demands for changing levels of energy consumption at intensely active synapses, at synapses that lack mitochondria, or at synapses in which mitochondria has been damaged. Therefore, the observed dynamic localization of glycolytic proteins to synapses may be essential to sustain changes in the activity of synapses in physiology and disease.

## Experimental Procedures

### Strains and genetics

Worms were raised on nematode growth media plates at 20°C using OP50 *Escherichia coli* as a food source (Brenner, 1974). For wild-type nematodes, *C. elegans* Bristol strain N2 was used. The following mutant strains were obtained through the *Caenorhabditis* Genetics Center: *juIs1[Punc-25::snb-1::gfp; lin-15(+)], jsIs682[Prab-3::gfp::rab-3], pfk-1.1(gk549413), pfk-1.1(gk758818), pfk-1.1(gk922689), pfkb-1.1(ok2733), gpd-3(ok2870), unc-57(ok310), unc-11(e47), unc-26(s1710), unc-13(e450), unc-13(e51), unc-104(e1265), klp-6(sy511), ric-7(nu447), syd-2(ju37), syd-2(ok217)*, and *unc-49(e407). aldo-1(tm5782)* and *pgk-1(tm5613)* were received from Shohei Mitani (Tokyo Women’s Medical University, Tokyo, Japan). *nuIs168[Pmyo-2::gfp; Punc-129::Venus::rab-3]* was provided by Jihong Bai (Fred Hutchinson Cancer Research Center, Seattle, Washington). *lev-8(ok1519)* was provided by Michael Koelle (Yale University, New Haven, Connecticut). *drp-1(tm408)* was provided by Marc Hammarlund (Yale University, New Haven, Connecticut). *syd-2(ju37); juIs330 [Punc-25::syd-2]* was provided by Mei Zhen (University of Toronto, Canada). Other strains used in the study are as follows: *olaIs1 [Ptph-1::mCherry; Ptph-1::cat-1::gfp], WyEx505 [Pttx-3::mCherry::erc; Pttx-3::GFP::rab-3]*, *oxSi91[Punc-17::ChIEF::mCherry]*, and *oxIs352 [Punc-47::ChR2::mCherry; lin-15(+)]*.

### Molecular biology and transgenic lines

Expression clones were generated using the Gateway system (Invitrogen). Detailed cloning information will be provided upon request. Transgenic strains (0.5–30 ng/μl) were generated using standard techniques (Mello and Fire, 1995) and coinjected with markers *Punc-122::gfp* or *Punc-122::rfp*. The following strains were generated: *olaEx1556 [WRM016aB04; Pttx-3::mCherry], olaEx2337 [Ppfk-1.1::pfk-1.1], olaEx1923 [Ppfk-1.1::pfk-1.1], olaEx1920 [Ptph-1::pfk-1.1], olaEx2085 [Ppfk-1.1::egfp], olaEx1641 [Ppfk-1.1::egfp; Ptph-1::mCherry], olaEx2014 [Ptph-1::pfk-1.1::egfp; Ptph-1:mCherry], olaEx2016 [Ptph-1::pfk-1.1::egfp; Ptph-1::mCherry::rab-3], olaEx2209 [Ptph-1::pfk-1.1::egfp; Ptph-1::mCherry::rab-3], olaEx2241 [Ptph-1::pfk-1.1::mCherry; Ptph-1::aldo-1::egfp], olaEx2249 [Ptph-1::pfk-1.1::mCherry; Ptph-1::gpd-3::egfp], olaEx2274 [Punc-47::pfk-1.1::egfp], olaEx2245 [Pttx-3:sl2:pfk-1.1:egfp*; *Pttx-3:mCh:rab-3]*, *olaEx2544 [Ptph-1::pfk-1.1G532E::egfp; Ptph-1::mCherry::rab-3], olaEx2545 [Ptph-1::pfk-1.1G532E]*, *olaEx2330 [Ptph-1::pfk-1.1::egfp::NLS; Ptph-1:mCherry:rab-3]*, *olaEx2290 [Ptph-1::pfk-1.1::egfp::NLS; Ptph-1:mCherry:rab-3]*, *olaEx2291 [Ptph-1::pfk-1.1::egfp::NLS; Ptph-1:mCherry:rab-3]*, and *olaex2671 [Ptph-1::mCherry::rab-3]*.

### Screen and positional cloning

Worms expressing CAT-1::GFP and cytosolic mCherry in NSM neuron (*olaIs1*) were mutagenized with ethyl methanesulfonate (EMS) as described previously (Brenner, 1974). The original screen was designed to identify mutants in genes required for synaptogenesis in the serotonergic neurons (NSM). The screen was performed as previously described (Colon-Ramos et al., 2007; Jin, 2005; Schaefer et al., 2000; Shao et al., 2013; Shen and Bargmann, 2003; Sieburth et al., 2005; Yeh et al., 2005). Briefly, F_1_ progeny of mutagenized P_0_ worms were cloned onto individual plates, and F_2_ progeny were screened using a Leica DM500B compound fluorescent microscope to visualize the distribution of the synaptic vesicular monoamine transporter (VMAT), CAT-1::GFP. NSM neurons were first inspected for appropriate cellular morphology and position, as well as axon and dendrite extension and arborization (by using the cytoplasmic mCherry marker). Next, the position, number, intensity, and distribution of the synaptic vesicle signal was assessed using our CAT-1::GFP marker. We screened through 2000 haploid genomes and identified 13 alleles with defects in synaptic patterning. These mutants fall into four categories: (1) Synaptic vesicle signal is diffusely distributed throughout neurites (6 mutants); (2) Synaptic vesicle signal is very dim or completely absent (4 mutants); (3) Synaptic varicosities appear larger than average (2 mutant); (4) Synaptic vesicle signal appears wild-type upon first examination; after extended exposure to hypoxic conditions under a coverslip (Pitts and Toombs, 2004) the synaptic vesicles became diffusely distributed. This fourth category is the mutant class *pfk-1.1(ola72)* belongs to.

The *ola72* allele was mapped to a 0.39Mbp region on chromosome X using single nucleotide polymorphisms and the CB4856 strain as described (Davis et al., 2005). Fourteen fosmids that cover this region were injected into *ola72* mutants and examined for rescue of synaptic vesicle clustering defects. *ola72/pfk-1.1* trans-heterozygotes were examined for complementation. Sanger sequencing was performed to identify the genetic lesion in the *ola72* allele.

### Cell autonomy and rescue of *ola72*

The *ola72* mutant phenotype was rescued by both the full *pfk-1.1* genomic sequence and *pfk-1.1* cDNA under the regulation of its endogenous promoter (fragment 1.5kb upstream of *pfk-1.1*). Cell-specific rescue in NSM was achieved with a plasmid driving the expression of *pfk-1.1* cDNA under the NSM-specific *tph-1* promoter fragment as described (Nelson and Colón-Ramos, 2013). *pfk-1.1* cDNA tagged with eGFP at its C-terminus under the NSM-specific promoter also rescued the *ola72* mutant phenotype (Figure S4H).

### Inducing hypoxia with glass coverslips and slides

Glass coverslips have been used to induce hypoxia in cell cultures (Pitts and Toombs, 2004). The reduced environment generated by mounting 10-15 live worms on glass slides was examined by using redox indicator resazurin (25ug/mL) dissolved in water. Resazurin when reduced to resorufin gives off fluorescence under yellow-green light (O’Brien et al., 2000). Using resazurin, we confirmed that worms between a glass coverslip and slide experience reduced, or hypoxic, conditions; Leica DM500B compound fluorescent microscope was used to acquire the images (Figures S1M and S1N). Gas permeable slides were made with Sylgard-184 (polydimethylsiloxane, or PDMS) (Dow Corning) according to manufacturer instructions.

### Inhibiting oxidative phosphorylation or glycolysis using a hypoxia chamber or pharmacological treatments

Hypoxia assays were performed using a chamber (similar to Billups-Rothernberg modular incubator chamber MIC-101) generously shared by Dr. Americo Esquibies (Yale University School of Medicine). Worms were mounted on a slide, placed in a hypoxia chamber and flushed for 4 minutes with nitrogen gas. The chamber was sealed and the worms were exposed to hypoxic conditions for 10 minutes. After the exposure, phenotypes were scored immediately under a Leica DM500B compound microscope. For carbon dioxide exposure, a smaller chamber was built using glass bottom culture dish (MatTek). Worms were placed in the culture dish and exposed to carbon dioxide for 10 minutes, and scored immediately. For pharmacological disruption of oxidative phosphorylation or glycolysis, worms were mounted on a slide with 10mM sodium azide (NaN_3_), 10uM of 2-deoxy-D-glucose (2-DG), or 1uM of oligomycin dissolved in muscimol, and scored after 10 minutes.

### Neuronal stimulation with pharmacological treatments or optogenetics

To pharmacologically stimulate GABA neurons, worms were mounted on a slide in 1mM levamisole (Sigma) and imaged immediately in spinning-disc confocal microscope (PerkinElmer Life and Analytical Sciences). As a control, 50mM of muscimol (Abcam) was used. To optogenetically stimulate GABA neurons, a strain expressing channelrhodopsin 2 in GABA neurons (*oxIs352*) (Liu et al., 2009) was used. All-trans-retinal and OP50-seeded plates were prepared as described previously (Liu et al., 2009). Briefly, 4uL of 100uM all-trans-retinal was added to 200uL of OP50 culture to seed each plate. Channelrhodopsin expressing worms were grown in all-trans-retinal seeded plate for minimum of 16 hours before stimulation. Worms were then mounted to a gas permeable PDMS slide set up, immobilized in muscimol, exposed to blue light for 5 minutes using a Leica DM500B compound microscope (approximately 0.6mW/mm^2^), and examined for formation of PFK-1.1 clusters. As a control, the same strains were grown without all-trans-retinal, and tested under the same conditions.

### Electrophysiology

For electrophysiology experiments, worms expressing the channelrhodopsin I/II chimera ChIEF in acetylcholine neurons (*oxSi91*) were dissected and exposed to 1uM oligomycin for 5 minutes. Patch-clamp physiology was conducted as previously described (Richmond et al., 1999; Richmond and Jorgensen, 1999). The stimulation protocol included a pre-pulse followed by a 5-second recovery period and 30-second stimulation train (10Hz). Test pulses were delivered at increasing time increments and normalized to the first spike in the train to assess recovery. Two test-pulses were delivered per animal at ≥15 seconds apart. By analyzing the overlap of the first and second pulses at intermediate time-points, we determined that a single test-pulse did not affect the amplitude of the second pulse. For repeated stimulus trains, the full protocol was repeated with 70 seconds between the end of the first train and the beginning of the following train.

### Behavioral experiments

*C. elegans* nematodes move in a wave-like fashion by executing dorsal-ventral body bends, and this sinusoidal body motion can be observed both on solid substrates and when worms are placed in an aqueous environment (Ghosh and Emmons, 2008). To examine how the locomotion of these animals is affected under hypoxic conditions, an Anaerobe Pouch System (Becton Dickinson & Co) was used to induce hypoxia. Animals were placed in M9 solution on slides, which were then placed inside the gas-impermeable pouch along with a wet paper towel and anaerobe gas-generating sachet. To examine how the locomotion of animals is affected under conditions of energetic stress, animals were placed in 10mM sodium azide (NaN_3_) solution dissolved in M9 on top of a slide. After 4 minutes of acclimation, the number of body bends was scored for 1 minute under a dissection scope. Bending of the head of the worm to one side and back to the initial side was counted as one body bend as illustrated in Figure 8H. M9 solution was used as a control and, as expected, no difference in the number of body bends between wild-type and *pfk-1.1* mutant animals were observed in this control (data not shown). *p values* were calculated using the Mann-Whitney *U* test.

### Microscopy, FRAP, and imaging

Images of fluorescently tagged fusion proteins were captured in live *C. elegans* nematodes using a 60 CFIPlan Apo VC, numerical aperture 1.4, oil-immersion objective on an UltraView VoX spinning-disc confocal microscope (PerkinElmer Life and Analytical Sciences). Worms were immobilized using 50mM muscimol (Abcam). Image J and Photoshop were used to analyze images, which were oriented anterior to the left and dorsal up. Maximum projections were used for all the confocal images, unless otherwise stated. Ratiometric images were generated by using the Volocity software (Perkin Elmer). For the Fluorescence Recovery After Photobleaching (FRAP) experiment, the spinning-disc confocal microscope and Volocity FRAP Plugin (Perkin Elmer) were used. Single plane images were acquired at 3-second intervals post-photobleaching. Acquired data was normalized using FRAP analysis in Volocity and fitted to a single exponential curve. For FRAP analysis, *p values* were calculated using the Mann-Whitney *U* test.

### Quantification of phenotypic expressivity

To quantify synaptic enrichment of the indicated proteins (synaptic vesicle proteins or PFK-1.1), fluorescence values for individual neurites (ventral neurite for the NSM neuron, Zone3 for the AIY neuron, and dorsal and ventral neurite for GABA neurons) were obtained through segmented line scans using ImageJ. A sliding window of 2μm was used to identify all the local fluorescence peak values and trough values for an individual neuron (the maximum and the minimum fluorescence values in a 2μm interval, respectively). Synaptic enrichment was then calculated as % Δ*F* /*F* as previously described (Bai et al., 2010; Dittman and Kaplan, 2006). Briefly, all the identified local maximum and minimum fluorescence values in a given neurite (local *F*_*peak*_ and local *F*_*trough*_) were averaged and used to calculate % Δ*F* /*F*, with % Δ*F* /*F* being the percent difference between average peak-to-trough fluorescence (*F*) defined as 100 × (*F*_*peak*_ – *F*_*trough*_)/*F*_*trough*_. All the images used in the quantification analyses were obtained using identical microscopy settings. For the quantification of the average number of PFK-1.1 puncta in AIY neurons, confocal images of AIY neurons were analyzed and scored for PFK-1.1 puncta in synaptic and asynaptic regions (Colon-Ramos et al., 2007; White et al., 1986). Statistical analyses were performed with Prism (GraphPad) and *p values* were calculated using the Mann-Whitney *U* test.

### Quantification of phenotypic penetrance

Animals were scored as displaying either “punctate” or “diffuse” phenotypes for synaptic vesicles proteins or PFK-1.1 after specified manipulations. “Punctate” or “diffuse” phenotypes were quantitatively defined in terms of their DF/F values. Briefly, we scored the distribution of synaptic vesicle proteins in wild-type or mutant animals by performing line scans, and calculated the DF/F for wild-type (N=16) and *pfk-1.1(ola72)* mutants (N=19). The average value for wild-type animals is 5.17 DF/F (fold increase of peak-to-trough fluorescence), while the average value for *pfk-1.1(ola72)* mutants is 2.94 DF/F fold. We then defined the “diffuse” distribution of vesicle clusters based on this average value observed for *pfk-1.1(ola72)* mutants and established a threshold at 3 DF/F for identifying an animal as either “diffuse” or “punctate”. Calculations, using this value, of the number of animals with a “diffuse” distribution of vesicle clusters reveal that approximately 20% of wild-type animals vs. 60% of *pfk-1.1(ola72)* mutant animals display a diffuse clustering phenotype under hypoxic conditions. We used these parameters as guidelines for scoring the phenotypic penetrance of all conditions and genotypes described in this study. We validated the approach by qualitatively scoring blindly the percentage of animals that exhibit a diffuse distribution of synaptic vesicle proteins for wild-type and for *pfk-1.1(ola72)* animals. We determined that the number we obtained after qualitative assessments of the phenotype and blind scoring were similar to the ones which were generated by measuring synaptic distribution of vesicular proteins using line scans, calculating the DF/F signal and then scoring individuals animals based on the 3 DF/F threshold definition of “diffuse”. Statistical analyses were performed with Prism (GraphPad) and *p values* were calculated using Fisher’s exact test.

### Quantification of colocalization

To quantify the relative distance between two fluorescently tagged proteins in the colocalization experiments, we used ImageJ and line scans of neurites to score the pixel fluorescence values in the green and red channels. The relative location corresponding to the maximum pixel fluorescence values for each channel was then used to calculate the distance between the two fluorescently tagged proteins.

## Author Contribution

J.N., S.J. and D.C-R. designed experiments. E.B. and E.J. performed the physiology experiments. J.N., G.T., K.U. and L.C. identified the allele of *pfk-1.1(ola72)* and the molecular lesion. J.N. and S.J. performed the cell autonomous rescue experiments. S. J. and L.R.-L. performed the subcellular localization studies, genetic and optogenetic analyses. S.J. performed all other experiments. S.J., J.N., L.R.-L. and D.C-R. analyzed and interpreted the data. S.J., J.N. and D.C.-R. wrote the paper.

## Acknowledgments

Cartoon in Figure 1A was modified with permission from the Neuron pages of WormAtlas (www.wormatlas.org) designed by Z.F. Altun and D.H. Hall.

We thank J. Bai, M. Zhen, J. Dittman, A. Esquibies, M. Hammarlund, M. Koelle, P. Lusk, R. Rawson, S. Margolis, S. Han, and members of the Colón-Ramos lab for strains, reagents and advice on the project. We thank N. Cook, J. Belina and M. Omar for technical assistance. Some strains were provided by the CGC, which is funded by NIH (P40 OD010440) and by the Mitani lab (Tokyo Women’s Medical University School of Medicine, Japan). Yuji Kohara (National Institute of Genetics, Japan) provided the *pfk-1.1* cDNA clone.

This work was funded by the following grants to D.C.-R. (R01 NS076558, a fellowship from the Klingenstein Foundation and the Alfred P. Sloan Foundation and a March of Dimes Research Grant) and to E.M.J (NIH R01 NS034307, NSF 0920069). S.J. was supported by the Cellular, Biochemical, and Molecular Sciences Predoctoral Training Program (T32 GM007223). E.M.J. is an Investigator of the Howard Hughes Medical Institute. J.N. was supported by a training grant 5 T32 NS 41228. L.R.L was supported by a diversity supplement to R01 NS076558. F.G.T. was supported by a fellowship from Universidad Ricardo Palma, Perú and the Research Experience for Peruvian Undergraduates (REPU) program.

